# A stem cell zoo uncovers intracellular scaling of developmental tempo across mammals

**DOI:** 10.1101/2022.10.13.512072

**Authors:** Jorge Lázaro, Maria Costanzo, Marina Sanaki-Matsumiya, Charles Girardot, Masafumi Hayashi, Katsuhiko Hayashi, Sebastian Diecke, Thomas B. Hildebrandt, Giovanna Lazzari, Jun Wu, Stoyan Petkov, Rüdiger Behr, Vikas Trivedi, Mitsuhiro Matsuda, Miki Ebisuya

**Author notes:** Correspondence to V. Trivedi, M. Matsuda or M. Ebisuya.

## Abstract

Differential speeds in biochemical reactions have been proposed to be responsible for the differences in developmental tempo between mice and humans. However, the underlying mechanism controlling the species-specific kinetics remains to be determined. Using *in vitro* differentiation of pluripotent stem cells, we recapitulated the segmentation clocks of diverse mammalian species varying in body weight and taxa: marmoset, rabbit, cattle and rhinoceros. Together with the mouse and human, the segmentation clock periods of the six species did not scale with the animal body weight, but were rather grouped according to phylogeny. The biochemical kinetics of the core clock gene HES7 displayed clear scaling with the species-specific segmentation clock period. However, the cellular metabolic rates did not show an evident correlation. Instead, genes involving biochemical reactions showed an expression pattern that scales with the segmentation clock period. Altogether, our stem cell zoo uncovered general scaling laws governing species-specific developmental tempo.*

## Introduction

Embryos from different mammalian species, despite using conserved molecular mechanisms, display differences in their developmental pace ^1,2^. For example, embryogenesis in humans consists of same series of events than in mice but takes 2-3 times longer ^3^. These proportional changes in the pace of development across species are known as developmental allochronies^4^.

A great example of developmental allochrony can be found during the segmentation of the vertebrate body axis. The pace of sequential formation of body segments is controlled by the segmentation clock, the oscillatory gene expression found in the cells of the pre-somitic mesoderm (PSM) ^5,6^. The oscillations of the segmentation clock are cell-autonomous, and their period differs across vertebrate species, with the human clock showing a period 2-3 times longer than the one of the mouse ^7–9^. Moreover, the segmentation clock is a convenient system to study interspecies differences in developmental pace as it can be modelled *in vitro* through the differentiation of pluripotent stem cells (PSCs) into PSM cells ^7,8,10,11^. Recently, *in vitro* recapitulation of the segmentation clock using mouse and human PSCs revealed that differences in the biochemical reaction speeds, including protein degradation rates and gene expression delays, are responsible for the interspecies differences in the mouse and human clocks ^12^. The protein degradation rate was also found to be associated to the species-specific pace of mouse and human motor neuron differentiation *in vitro* ^13^. However, whether this mechanism constitutes a general principle of development, as well as the underlying cause behind the interspecies differences in biochemical reaction speeds remains unknown. This is in part because all studies to date have been limited to mouse and human comparisons, making it challenging to examine general relationships between developmental tempo and other cellular parameters.

With the recent expansion of the PSC technologies, we can now extend our knowledge of development outside of the classical mouse and human models. *In vitro* differentiation of PSCs from different mammalian species can be used to recapitulate key features of development and study them under similar experimental conditions ^14^. For example, comparisons of human and primate PSC-derived brain models have helped reveal unique properties of human brain development ^15,16^. Thus, in *vitro* models of development from multiple species represent a great opportunity to perform interspecies comparisons of cell- and tissue-autonomous processes. In this work, we recapitulated *in vitro* the segmentation clock of four novel mammalian species in addition to the mouse and human. We then used this “stem cell zoo” platform to systematically investigate the general mechanism behind developmental allochrony.

## Results

### A stem cell zoo platform to study interspecies differences in the segmentation clock

Using the segmentation clock as a model, we sought to expand previous results in the mouse and human by establishing a general platform to study differences in developmental tempo across multiple mammalian species (Fig. 1A). First, we collected embryonic stem cells (ESCs) and induced pluripotent stem cells (iPSCs) from diverse mammalian species, including common marmoset (*Callithrix jacchus*), rabbit (*Oryctolagus cuniculus*), cattle (*Bos taurus*) and southern white rhinoceros (*Ceratotherium simum*). Together with mouse and human PSCs, these species show adult body weights spanning from 50 grams to 2 tonnes, and gestation lengths ranging from 20 days to 17 months. Given the wide range of body weights and gestation lengths in these six species, we would expect significant differences in their developmental tempo. Moreover, these species belong to three distinct phylogenetic clades: Primates (marmoset and human), Glires (mouse and rabbit) and Ungulates (cattle and rhinoceros), constituting a diverse sampling of mammalian species unprecedented for developmental studies. We used rabbit ESCs ^17^, cattle ESCs ^18^, rhinoceros ESCs ^19^ and marmoset iPSCs ^20^ to induce PSM-like cells from these species following protocols already described for mouse epiblast stem cells (EpiSCs) and human iPSCs (Fig. S1) ^7,12^. We used identical culture conditions when measuring the induced PSM cells, minimizing the effect of external factors on our quantifications. After 2-3 days of induction, cells showed mesoderm-like morphology and expression of the PSM fate marker TBX6 (Fig. 1B and C). The efficiency of differentiation, measured by TBX6 expression, was around 80-95% in all species (Fig. 1C and Fig. S2A, B). All subsequent measurements were done on the most efficient day of differentiation for each species. To further characterize the identity of these PSM-like cells and compare them to the previously described mouse and human PSM, we performed bulk RNA-sequencing (RNA-seq) on PSCs and PSM-like cells of the six species. This confirmed the general expression of PSM markers as well as a similar anterior-posterior identity across species, with all the induced PSM cells showing a thoracic-lumbar fate (Fig. S3A and B). The induced PSM-like cells are hereafter referred to as PSM.

**Figure 1:**
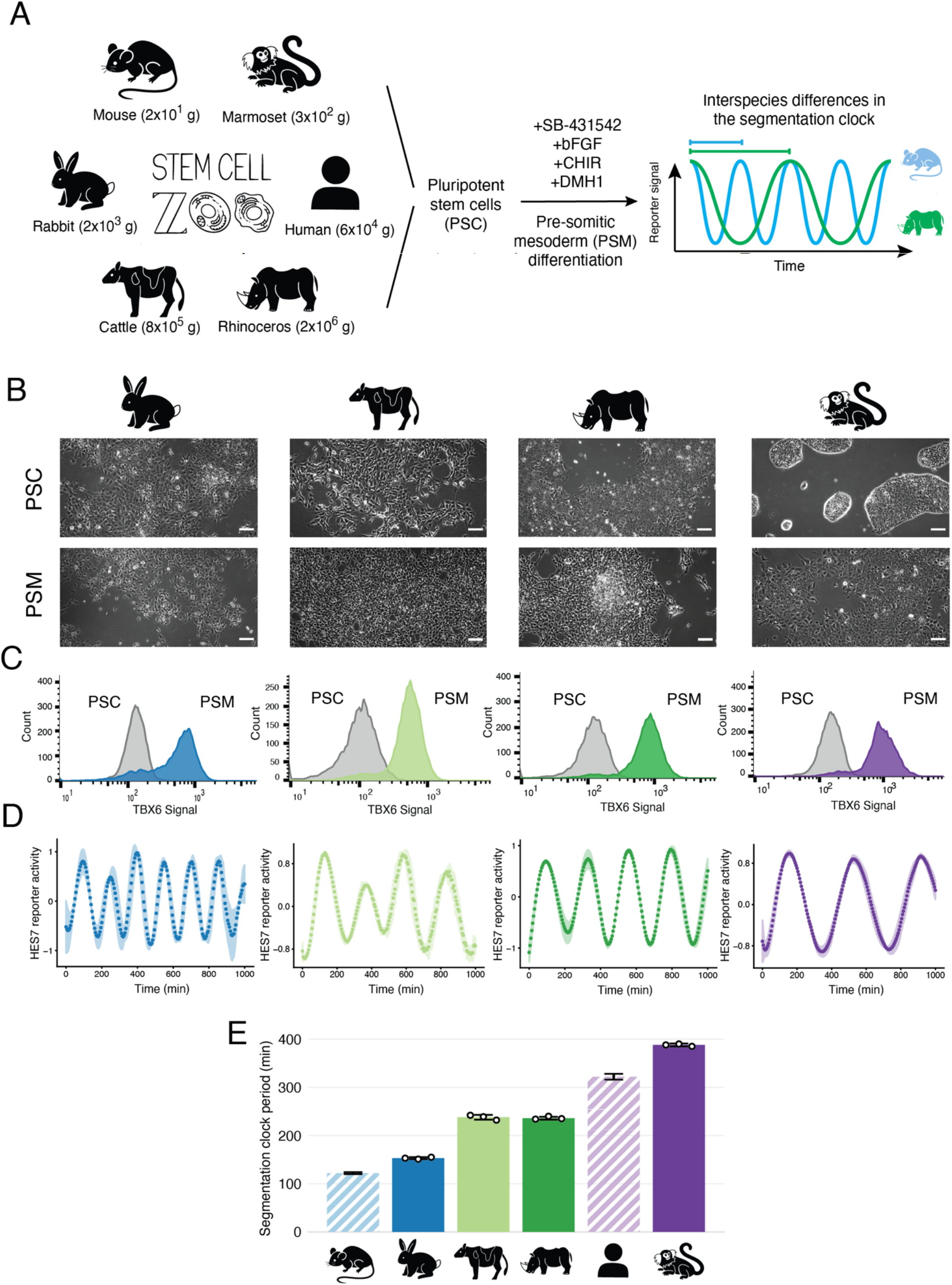
Recapitulation of the segmentation clock using stem cells from diverse mammalian species (A) Schematic illustration of the differentiation of mammalian PSCs towards PSM. Cells differentiated under the same conditions show species-specific segmentation clock periods. Average adult body weight of each species is displayed. (B) Bright-field images of PSCs and PSM cells from each species. Scale bars are 100 microns. (C) Representative histogram of flow cytometry analysis of a PSM marker TBX6 in PSCs (grey) and PSM cells (coloured) of each species. (D) Normalised HES7 reporter activity in PSM cells of each species. Shading indicates indicate means ± SD (n = 3). The signal has been detrended and amplitude-normalised using pyBOAT (see methods). (E) Oscillatory periods estimated from Fig. S4. Error bars indicate means ± SD. (n= 3). Human and mouse data (striped bars) are extracted from Matsuda et al., Science, 2020.

To visualize the oscillations of the segmentation clock, we introduced a HES7 promoter luciferase reporter, which displayed clear oscillations of HES7 expression in all species. In mammals, HES7 constitutes the core oscillatory gene of the segmentation clock. Rabbit PSM oscillated with a period of 153 ± 2 min (mean ± SD), followed by cattle PSM with a period of 238 ± 5 min, rhinoceros PSM with a period of 236 ± 3 min, and finally marmoset PSM exhibited the longest period of 388 ± 3 min (Fig. 1D and S4). Except for the rhinoceros, which lacks embryological data, the ranking order of the measured periods in the different species corresponds to the order of their roughly estimated *in vivo* somite formation periods (Table 1) ^21–23^. For example, the marmoset, which presents the longest period, is known to have a particularly slow pace of early embryo development ^24^. These results demonstrate that PSCs from different mammalian species can be used to recapitulate the segmentation clock *in vitro*, with the PSM of each species oscillating at a defined period. Together with the previous data of the mouse and human periods (122 ± 2 min and 322 ± 6 min, respectively) ^12^, the segmentation clocks of *in vitro* differentiated PSM cells showed almost a 4-fold difference from the fastest to the slowest species (Fig. 1E). Thus, our stem cell zoo serves as an ideal platform to investigate the cause of interspecies differences in the segmentation clock period as well as to determine whether there is any general relationship between developmental tempo and the organism characteristics.

**Table 1:** Estimation of the *in vivo* somite formation period The approximate period of *in vivo* somite formation was calculated using studies describing the number of somites in staged embryos.

| Rabbit |  |  | Cattle |  |  | Marmoset |  |  |
| --- | --- | --- | --- | --- | --- | --- | --- | --- |
| Embryonic day | nº somites | nº embryos | Embryonic day | nº somites | nº embryos | Embryonic day | nº somites | nº embryos |
| 10 | 26,5 | 40 | 23 | 16-22 | Not disclosed | 50 | 8 | 3 |
| 11 | 37,4 | 61 | 24 | 23-27 |  | 60 | 26,5 | 2 |
| 12 | 46,6 | 57 | 25 | 28-32 |  |  |  |  |
|  |  |  | 26 | 33-36 |  |  |  |  |
|  |  |  | 27 | 37-40 |  |  |  |  |
|  |  |  | 28 | 41-44 |  |  |  |  |
| Estimated period of in vivo somite formation (min) | Reference |  | Estimated period of in vivo somite formation (min) | Reference |  | Estimated period of in vivo somite formation (min) | Reference |  |
| 143 | Naya et al. 1991 <sup>21</sup> |  | 310 | Haldiman 1980 <sup>23</sup> |  | 778 | Chambers et al. 1985 <sup>22</sup> |  |

### The segmentation clock period does not scale with adult body weight but scales with embryogenesis length

The gestation length, the metabolic rate, as well as many other bodily parameters, are known to scale with the animal body weight ^25–27^. Larger species tend to have a longer gestation and a slower metabolism. We thus hypothesized that the observed differences in the segmentation clock period could be related to body weight. However, no correlation between the average adult body weight of each species and its segmentation clock period could be found (Fig. 2A). Similarly, the gestation length did not correlate with the segmentation clock period (Fig. 2B). We then checked general hallmarks of development and found that the embryogenesis length, defined as the time going from fertilization to the end of organogenesis (i.e., the last Carnegie stage when the secondary palate fuses), correlated highly with the segmentation clock period (Fig. 2C). This suggests that the segmentation clock can serve as a good proxy for quantifying embryonic developmental tempo and that the pace and overall length of early development are tightly connected. Furthermore, the three distinct phylogenetic clades, Primates, Glires and Ungulates, corresponded to slow, fast and intermediate segmentation clock periods respectively, pointing to a relation between developmental tempo and evolutionary distance (Fig. 2D).

**Figure 2:**
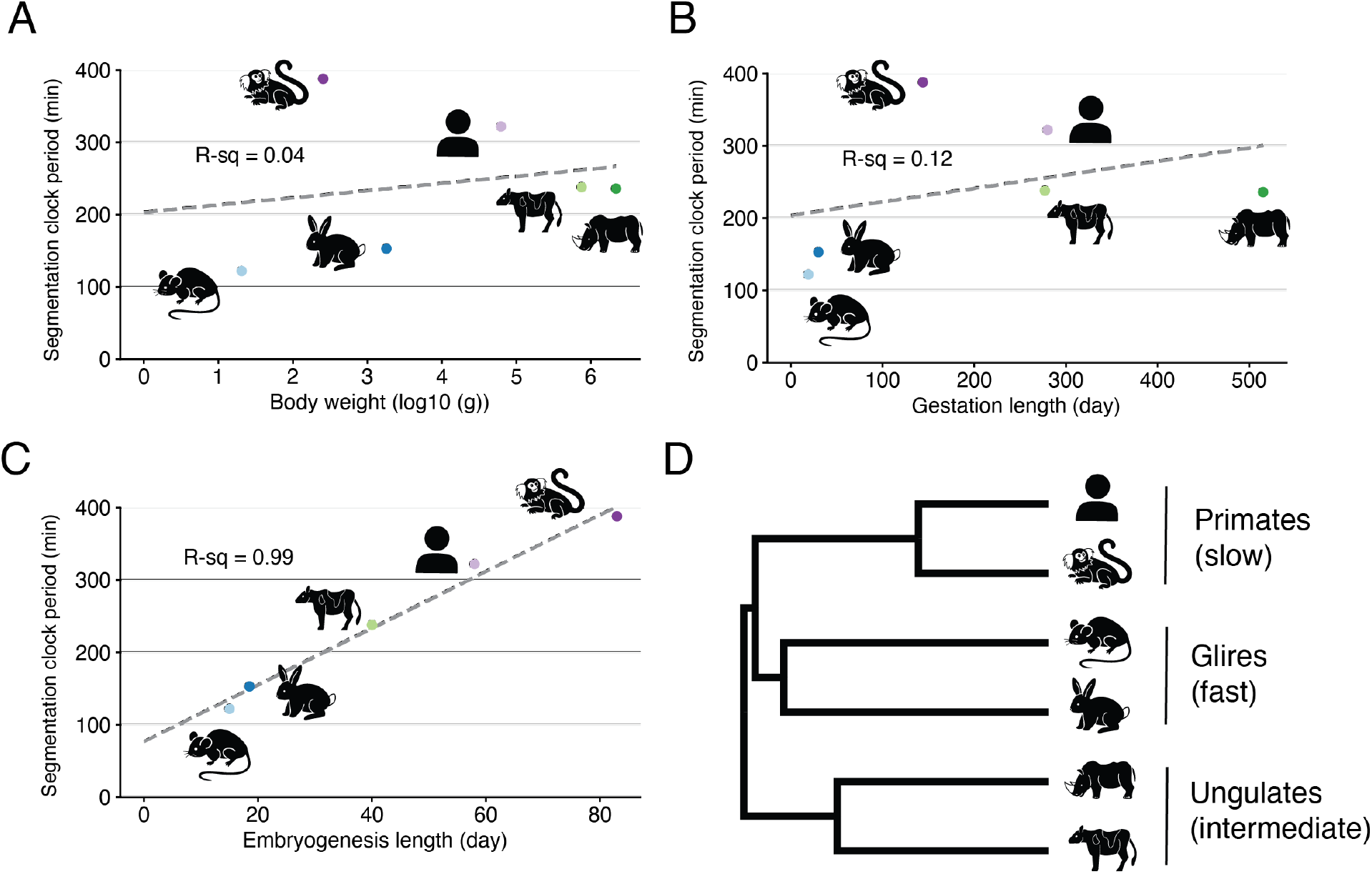
Correlations between the segmentation clock period and the animal characteristics (A) Scatterplot showing the relationship between the log10 average adult body weight and the segmentation clock period. (B) Scatterplot showing the relationship between the gestation length and the segmentation clock period. (C) Scatterplot showing the relationship between the embryogenesis length and the segmentation clock period. The rhinoceros is missing as it lacks embryological data. (A-C) Dashed lines represents linear fitting. R-squared values are shown. Values of the animal characteristics can be found in Table 2 (D) Phylogenetic tree of the six species used in this study. This tree represents a subset of the complete mammalian tree (see methods). Names of the common clades are shown.

### Biochemical reaction speeds scale with the segmentation clock period

The speed of biochemical reactions has been shown to change with developmental tempo ^12,13^. Human PSM shows slower degradation of HES7 mRNA and protein as well as longer delays in HES7 gene expression as compared to mouse PSM ^12^. To determine whether this is a general trend, we measured the degradation rates and delays affecting the regulatory negative feedback loop of HES7 in our different species (Fig. 3A). We focused on the HES7 protein degradation rate and the delay in transcript processing caused by HES7 introns, as these parameters were shown to be the most relevant for the period difference between the mouse and human segmentation clocks ^12^. First, we measured the HES7 protein degradation by overexpressing the human HES7 sequence fused with a luciferase reporter and then halting its expression with doxycycline (Fig. 3B). The human HES7 sequence was used for measurements in all species since it has been demonstrated that human and mouse HES7 sequences can be interchanged without affecting the pace of the clock or the HES7 reaction kinetics ^12^. The observed half-lives of HES7 protein were 24 ± 0.8 min, 33 ± 2 min, 32 ± 2 min and 46 ± 3 min in rabbit, cattle, rhinoceros and marmoset PSM, respectively (Fig. 3D and S5A). Together with the previously reported values in mouse and human PSM ^12^, the HES7 protein degradation rate was highly correlated with the segmentation clock period, with slower species showing longer HES7 half-lives (Fig. 3F). We then measured the delay caused by HES7 introns, by using HES7 promoter-luciferase reporters with and without HES7 intron sequences and estimating the phase difference between oscillations from both reporters (Fig. 3C). The intron delays were found to be 24 ± 3 min, 37 ± 2 min, 36 ± 3 min and 54 ± 0 min in rabbit, cattle, rhinoceros and marmoset PSM, respectively (Fig. 3E). Similar to the protein half-life, HES7 intron delay was highly correlated with the segmentation clock period (Fig. 3F). Note that protein degradation and intron delay are two completely different biological processes and that their contribution to the HES7 regulatory feedback loop can theoretically vary independently across species. For example, a species could alter its segmentation clock period by mostly modifying the HES7 protein half-life but not the intron delay. Nevertheless, all six species change their protein half-life and intron delay proportionally, indicating that species-specific degradation and intron delay may be co-regulated. Taken together, we found that the speed of biochemical reactions varies across species, and that these differences correlate very well with the segmentation clock period. This indicates that changes in the biochemical rates might be a general mechanism to control developmental tempo.

**Figure 3:**
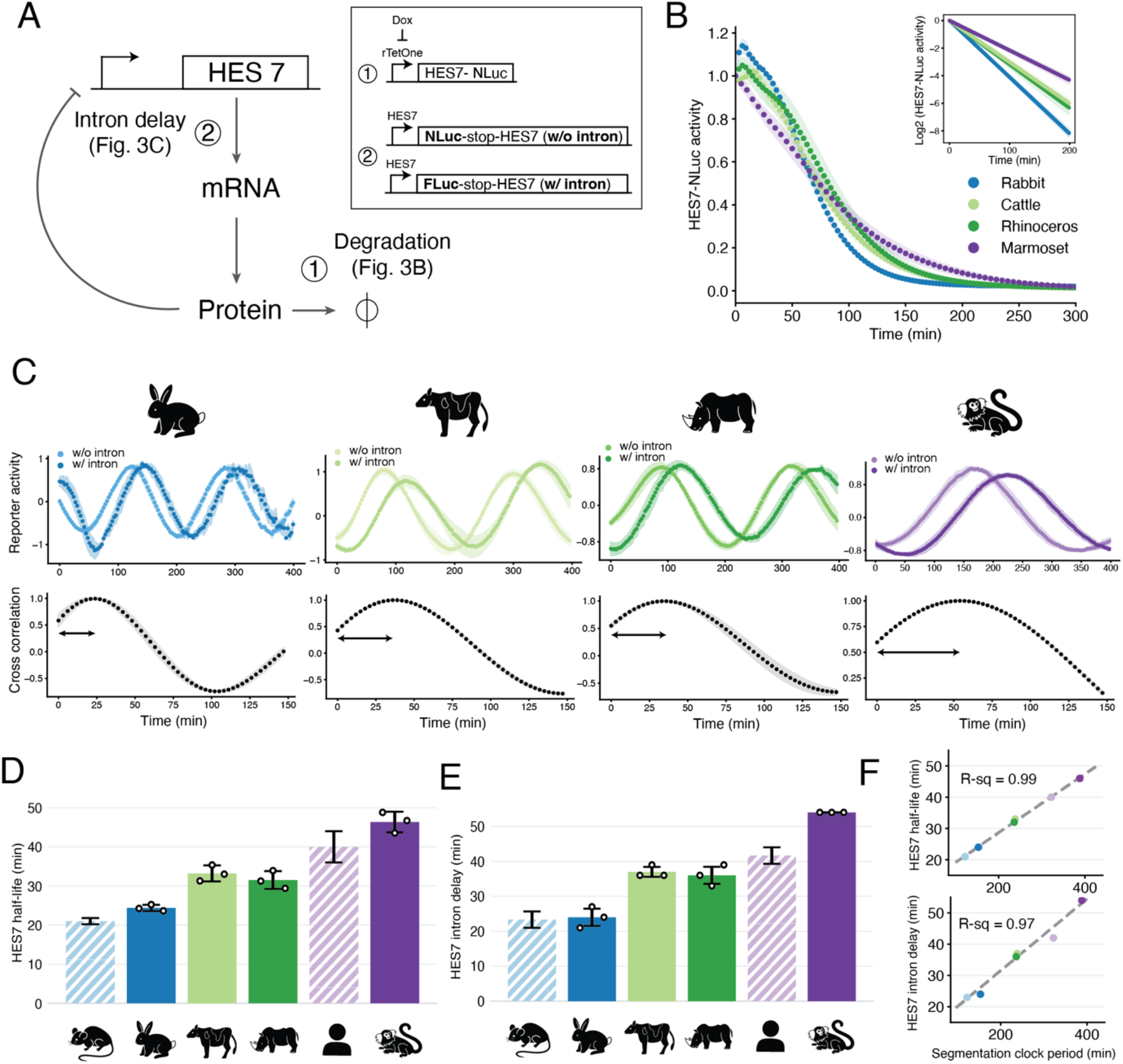
Measuring biochemical parameters of HES7 (A) Schematic representation of the delayed negative feedback loop of HES7. Protein degradation and intron delay were measured in the indicated panels. Reporters used for these two assays are shown. NLuc: NanoLuc, Fluc: Firefly Luciferase. (B) HES protein degradation assay. The transcription of a HES7 protein fused with NLuc was halted upon the addition of doxycycline (Dox). The signal decay of Nluc was monitored. Inset represents the slope of the fitted lines used to quantify the protein half-life. (C) HES7 intron delay assay. Reporter constructs without (w/o) and with (w/) HES7 introns were measured simultaneously (top). The cross-correlation of the two reporters was calculated (bottom). (B and C) Shading indicates means ± SD (n = 3). (D) HES7 protein half-lives estimated from Fig. S5A. (E) HES7 intron delays estimated from (C). (D and E) Error bars indicate means ± SD. Human and mouse data (striped bars) are extracted from Matsuda et al., Science, 2020. (F) Scatterplot showing the relationship between the segmentation clock period and the measured biochemical parameters (top: HES7 protein half-life, bottom: HES7 intron delay). Colour scheme representing species is the same as (D) and (E). Dashed lines represent linear fitting. R-squared values are shown.

### Metabolic rates do not directly scale with the segmentation clock period

Differences in metabolism have been proposed as a potential mechanism underlying the interspecies differences in biochemical reaction speeds and therefore the species-specific segmentation clock period. A recent report showed that mouse PSM cells, with a fast period, have higher mass-specific metabolic rates than human PSM ^28^. For this reason, we sought to examine the relationship between the segmentation clock period and the cellular metabolic rate using our stem cell zoo. To normalise our metabolic measurements to cellular size, we first measured the cellular volume of PSM cells (Fig. 4A). While different species showed differential cell volumes, we observed poor scaling with the segmentation clock (R-sq = 0.46; Fig. 4B and S6A). We then used the Seahorse analyser to measure the basal oxygen consumption rate (OCR), an indicator for mitochondrial respiration, across species (Fig. 4C). We focused on the mouse, rhinoceros, human and marmoset as they are the species covering a wider range of body weights and segmentation clock periods. Cell volume-specific OCR values were found to be different across species, with mouse PSM having a faster metabolic rate than human PSM, as previously reported ^28^. However, both rhinoceros and marmoset PSM showed faster respiratory rates than mouse, suggesting no correlation between the OCR and the segmentation clock period (Fig. 4D). OCR before cell volume normalisation, despite presenting a slightly different trend, also showed poor scaling with the segmentation clock (Fig. S6B). Next, we performed the real-time ATP rate assay to assess potential differences in the glycolytic and mitochondrial function (Fig. 4C). While the total ATP levels were similar across species (Fig. 4E), the origin of ATPs differed. Mouse PSM was the most glycolytic, showing the highest cell volume-specific glycolytic rate, followed by marmoset, human and finally rhinoceros PSM (Fig. 4F and S6F). This evidences a lack of scaling between the glycolytic rate and the segmentation clock period. Mitochondrially produced ATP showed an opposite trend, similar to the one of OCR (Fig. 4G). ATP production rates before cell volume normalisation also showed no correlation with the segmentation clock period (Fig. S6C to E). Collectively, these results suggest that metabolic rates, despite being different across species, do not directly scale with the species-specific segmentation clock period.

**Figure 4:**
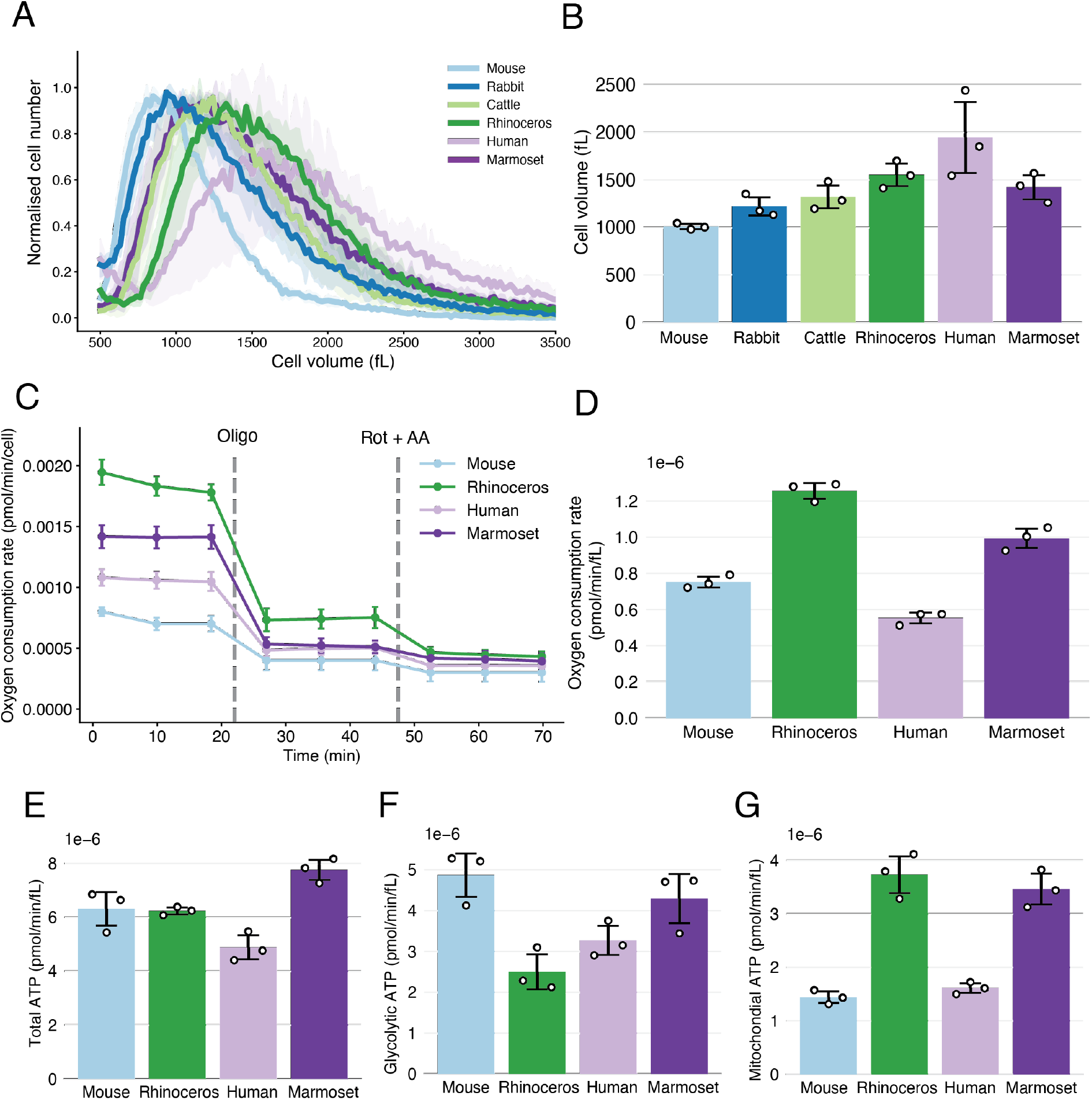
Measuring cellular metabolic rates (A) Histogram showing the size distribution of PSM cells of each species. Total cell number was normalised. Shading indicates means ± SD (n = 3). (B) Median cell sizes estimated from (A). (C) Oxygen consumption rate per cell measured over the course of the Seahorse real-time ATP rate assay in PSM cells of each species. Oligomycin (Oligo) and Rotenone + Antimycin A (Rot + AA) were added at the marked time points. Error bars indicate mean ± SD. (D) Volume-specific oxygen consumption rate across species. (E) Volume-specific ATP production rate across species. (F) Volume-specific glycolytic rate of ATP production across species. This measurement is equivalent to the glycolic proton efflux rate as per the stoichiometry of the glycolysis reaction. (G) Volume-specific mitochondrial rate of ATP production across species. (B, D-G) Error bars indicate means ± SD (n = 3).

### A transcriptional profile that scales with the segmentation clock period

After observing the disconnection between the segmentation clock period and cellular metabolism, we hypothesized that the pace of development could be controlled by species-specific gene expression profiles. Therefore, we explored our RNA-seq data to characterize the potential transcriptomic signatures of developmental allochrony. For this, we compared the expression levels of more than 10000 orthologous protein-coding genes across our six species. Principal component analysis (PCA) revealed that samples mostly cluster by species instead of tissue (Fig 5A and Fig. S7A). Similarly, hierarchical clustering of the RNA-seq data showed that samples preferentially cluster by species (Fig. S7B). This is in contrast with many observations made in adult tissues, where cell types cluster before species ^29^, but could be explained by considering that PSM cells are a relatively early cell type which remains transcriptionally close to PSCs. Interestingly, the first PCA axis clustered species by their segmentation clock period, grouping the fast (rabbit and mouse), intermediate (cattle and rhinoceros) and slow (marmoset and human) species together in ascending order (Fig. 5A). This suggests that differential gene expression profiles could be related to the species-specific segmentation clock periods. We further identified genes correlating with developmental time by calculating the Spearman correlation coefficient between the gene expression level in the PSM and the segmentation clock period for the six species (Fig. 5B and S8A and B). Gene set enrichment analysis (GSEA) demonstrated that the negatively correlated genes, which are genes expressed higher in faster species, showed enrichment in gene ontology (GO) terms of mRNA processing and protein catabolism (Fig. 5C). This, together with our previous results indicating that HES7 biochemical reactions are accelerated in species with faster segmentation clock periods, suggests that the speed of biochemical reactions could be controlled transcriptionally. Visualization of the enriched GO terms revealed they are highly interconnected, supporting the idea that these processes are regulated in a coordinated manner (Fig. 5D). Other basic biochemical processes, such as mRNA splicing, transcription elongation and nuclear transport, were also enriched in negatively correlated genes (Fig. 5D). Similar results could be obtained when ranking the genes expressed in PSCs (Fig. S8B). In contrast, enriched terms in positively correlated genes formed a much smaller cluster, indicating that few biological processes correlate positively with the segmentation clock period (Fig. 5D). Together, these results show that species with faster segmentation clock periods present higher expression of genes related to biochemical reactions. This suggests that the species-specific segmentation clock periods might be derived from species-specific gene expression profiles controlling basic biochemical processes (Fig. 5E).

**Figure 5:**
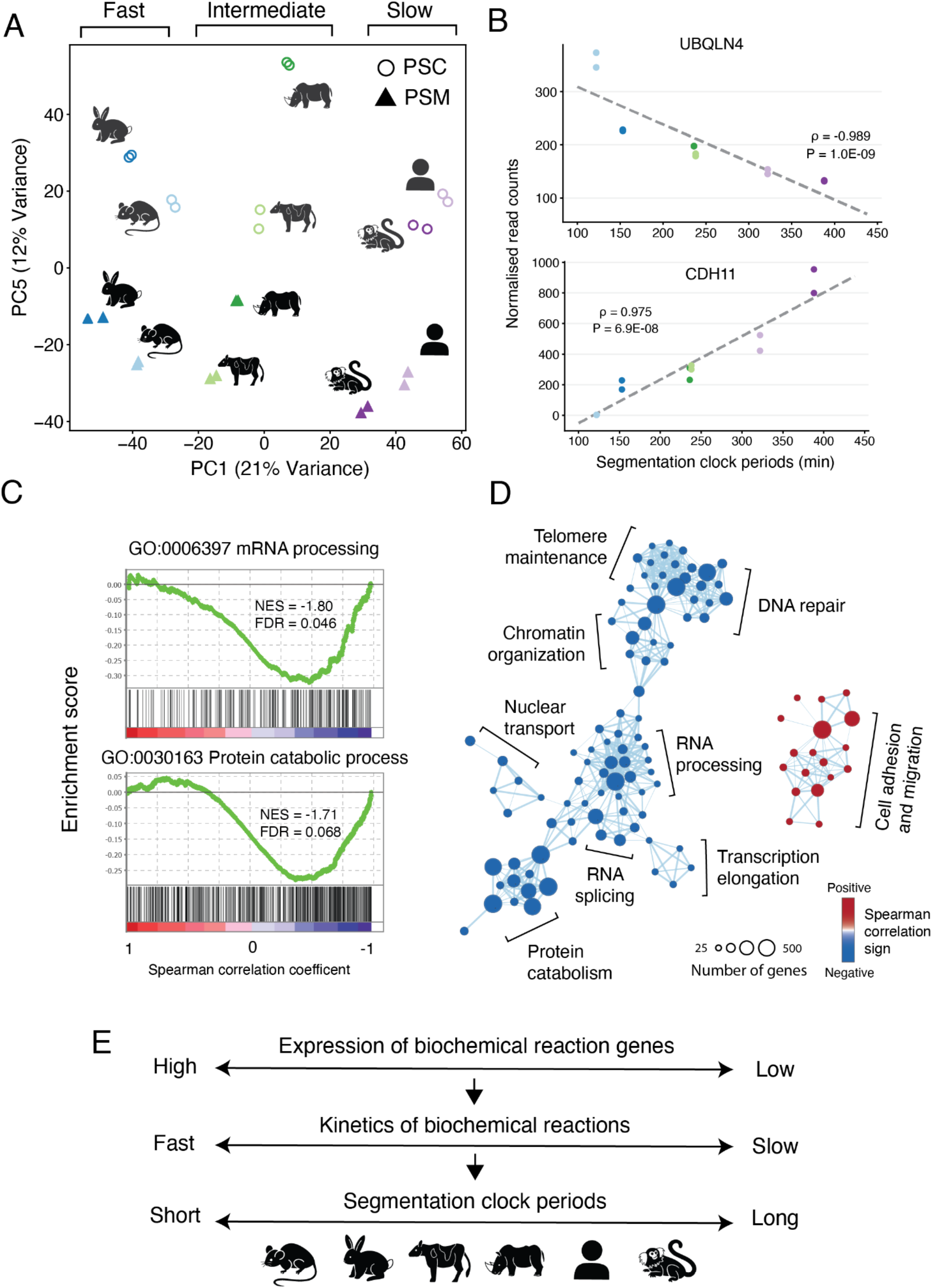
The transcriptomic signature of species-specific developmental tempo (A) Principal component analysis (PCA) of the log-transformed Gene length corrected trimmed mean of M-values (GeTMM) normalised reads in the bulk RNA-seq. Two biological replicates of PSCs (circles) and PSM cells (triangles) of six species were used. Components 1 and 5 are shown. The variance explained by each component is indicated. (B) Scatterplots showing the relationship between the GeTMM-normalised gene expression levels in PSM cells and the segmentation clock period across species. Colour scheme representing species is the same as (A). Spearman correlation coefficients (ρ) and p-values are shown in the plots. Example genes with top negative and positive ρ values are displayed. (C) Gene set enrichment analysis (GSEA) for two gene ontology (GO) terms: mRNA processing and Protein catabolism processes. These terms are enriched in genes negatively correlated with the segmentation clock period in PSM samples. Normalised Enrichment Score (NES) and False Discovery Rate (FDR) are displayed. (D) Enrichment map network of genes that showed correlated expression with the segmentation clock period. Each dot represents an enriched GO biological process term. Two terms are connected if they have a high overlap of genes. Related functional terms tend to cluster together. Circle size represents the number of genes in that process. Blue and red colours represent processes correlating negatively and positively with the segmentation clock period, respectively. (E) Proposed scheme from this study.

## Discussion

We have established a unique experimental platform, the stem cell zoo, that allowed us to explore which cellular parameters or animal properties correlate with developmental tempo. To this end, we successfully recapitulated *in vitro* the segmentation clock of four novel mammalian species: rabbit, cattle, rhinoceros and marmoset. By expanding the classic human and mouse models, we have revealed the existence of a general scaling law between developmental tempo and the speed of biochemical reactions. We further found that genes related to biochemical processes show an expression pattern that negatively correlates with the segmentation clock period, providing evidence of the potential transcriptional regulation of developmental allochrony. Altogether, quantitative measurements of the *in vitro* segmentation clock in six different mammalian species enabled us to establish strong correlations which would have otherwise remained elusive.

One potential mechanism underlying the differences in developmental tempo has been proposed to be metabolism. Recent studies done in mouse and human cells described the impact of metabolism on the segmentation clock or other developmental processes such as corticogenesis ^28,30^. While it is clear that changes in metabolism can influence the developmental rate within a given species, the metabolic characterization done in this study indicates that intrinsic species-specific differences in the segmentation clock period or the speed of HES7 biochemical reactions are not directly derived from differential metabolic rates. So far, the ultimate mechanism by which some species display slower or faster biochemical reactions remains unknown. However, this study provides a list of biochemical process genes that correlate with segmentation clock periods as a concrete clue. These genes could be exploited in future studies to further understand and even manipulate the tempo of the segmentation clock in different species.

The stem cell zoo also revealed that while the segmentation clock period does not scale with the adult body weight, is it highly correlated with the length of embryogenesis. This suggests that studying the species-specific segmentation clock periods may lead to a better understanding of how the embryogenesis pace and duration are established across species. However, more species covering a wider range of phylogenies and different cell types are necessary to confirm whether the reported findings could constitute a universal principle of mammalian development. The expansion of the stem cell zoo platform will be useful to study further interspecies differences.

## Acknowledgements

This work was supported by EMBL; the European Research Council (ERC) under the European Union’s Horizon 2020 research and innovation program (grant agreement No. 101002564) (to M.E.); Takeda Science Foundation (to M.E.); Project PGC2018-097872-A-I00 (MCIU/AEI/FEDER, UE) funded by the Spanish Ministry of Science, Innovation and Universities (MCIU) and co-funded by the European Regional Development Fund (ERDF, EU) (to M.E.); PRESTO (grant number JP20332265) from Japan Science and Technology Agency (JST) (to M.M.); the Boehringer Ingelheim Fonds (BIF) PhD fellowship (to J.L.); and the Japan Society for the Promotion of Science (JSPS) fellowship (to M.S-M.); the Deutsches Zentrum für Herz-Kreislauf-Forschung (grant number 81Z0300201) (to R.B.). J.W. is a New York Stem Cell Foundation–Robertson Investigator and Virginia Murchison Linthicum Scholar in Medical Research.

The human, mouse and rabbit PSC lines were provided by the RIKEN BRC through the National BioResource Project of the MEXT, Japan.

We are thankful to the members of the Ebisuya lab for advice and critical feedback during the whole duration of the project; to Júlia Charles Aymamí for designing the animal icons and stem cell zoo logo; to Jaroslaw Sochacki for advice on the culture of stem cells; to the BIF network for the feedback during the progress meetings; to the Flow Cytometry Unit of the Center for Genomic Regulation for consultation in data acquisition and analysis; to Jelle Scholtalbers from the Genome Biology Computational Support for support with the EMBL Galaxy platform; to Fumio Nakaki for advice in RNA-seq data analysis; to Joaquina Delas Vives and Teresa Rayon for experimental advice; to Gregor Mönke for implementing features into pyBOAT that facilitated the analysis of the data; to Susanne Holtze, Frank Göritz and Robert Hermes from the Leibniz Institute for Zoo and Wildlife Research for particiating on the rhinoceros semen and oocyte collections necessary to produce the rhinoceros ESCs line; to Michal Zaluski and his staff for their support during the rhinoceros oocyte collection at the Silesian Zoological Garden in Chorzow; to Miriam Wiesner and her staff for the participation during the rhinoceros semen collection at the Salzburg Zoo; to Cesare Galli and Silvia Colleoni for their input during the in vitro production of the rhinoceros embryos; to Juan Carlos Izpisua Belmonte for facilitating the obtention of cattle ESCs; to Kyoto University for facilitating the obtention of the human iPSCs; to Arata Honda for depositing the rabbit ESCs in the RIKEN BRC, to Milena Marinkovic, Carina Vibe, Guillermo Martínez Ara, Marc Duque Ramírez, James Briscoe and Teresa Rayon for their comments on the manuscript.

## Author contributions

M.M. and M.E. conceived the project. J.L. performed most of the experiments and analysed the data. M.M. provided advice and performed experiments with human and mouse PSCs. M.C. optimized cattle cell culture and performed experiments with cattle ESCs. M.S-M. optimized cattle cell culture and performed the HES7 protein degradation assays of cattle cells. C.G. performed gene expression analyses from the raw bulk RNA-seq data and assembled the multi-species gene ortholog table. M.H and K.H provided a table of orthologous genes between the human, mouse and rhinoceros. S.D, T.B.H. and G.L. provided the rhinoceros ESCs and advice to culture them. J.W. provided the cattle ESCs and advice to culture them. S.P. and R.B provided the marmoset iPSCs and advice to culture them. J.L and M.E wrote the manuscript with feedback from all authors. M.E., M.M. and V. T. supervised the project.

## Supplementary figures

**Figure S1:**
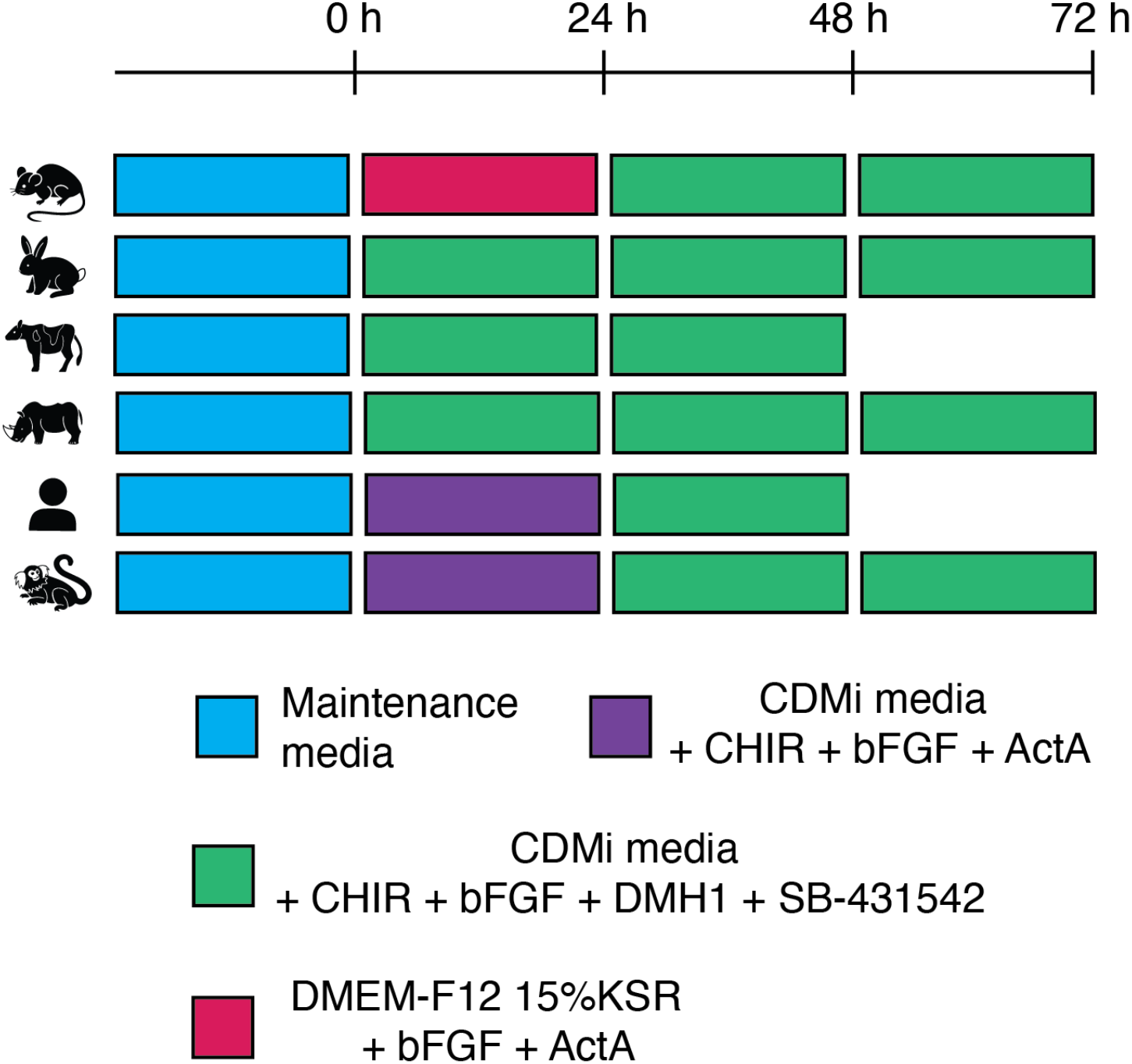
Induction protocols of PSM-like cells Schematic representation of the protocols used to induce PSM-like cells from PSCs of different mammalian species. PSCs were cultured in the maintenance media optimized for individual species, and the media were changed to the differentiation media at time 0. The differentiation protocol consists of culturing the cells for 2 to 3 days (depending on the species) in CDMi media containing Chiron (CHIR, WNT signalling activator), bFGF, DMH1 (BMP signalling inhibitor) and SB-431542 (TGF-beta signalling inhibitor). Prior to that, Human and Marmoset cells were first cultured for 24 hours in CDMi media with CHIR, bFGF and Activin A. Mouse cells were first cultured for 24 hours in their maintenance media without IWR-1. PSM-like cells of all species were analysed under the same culturing conditions during the most efficient day of differentiation (see Fig. S2).

**Figure S2:**
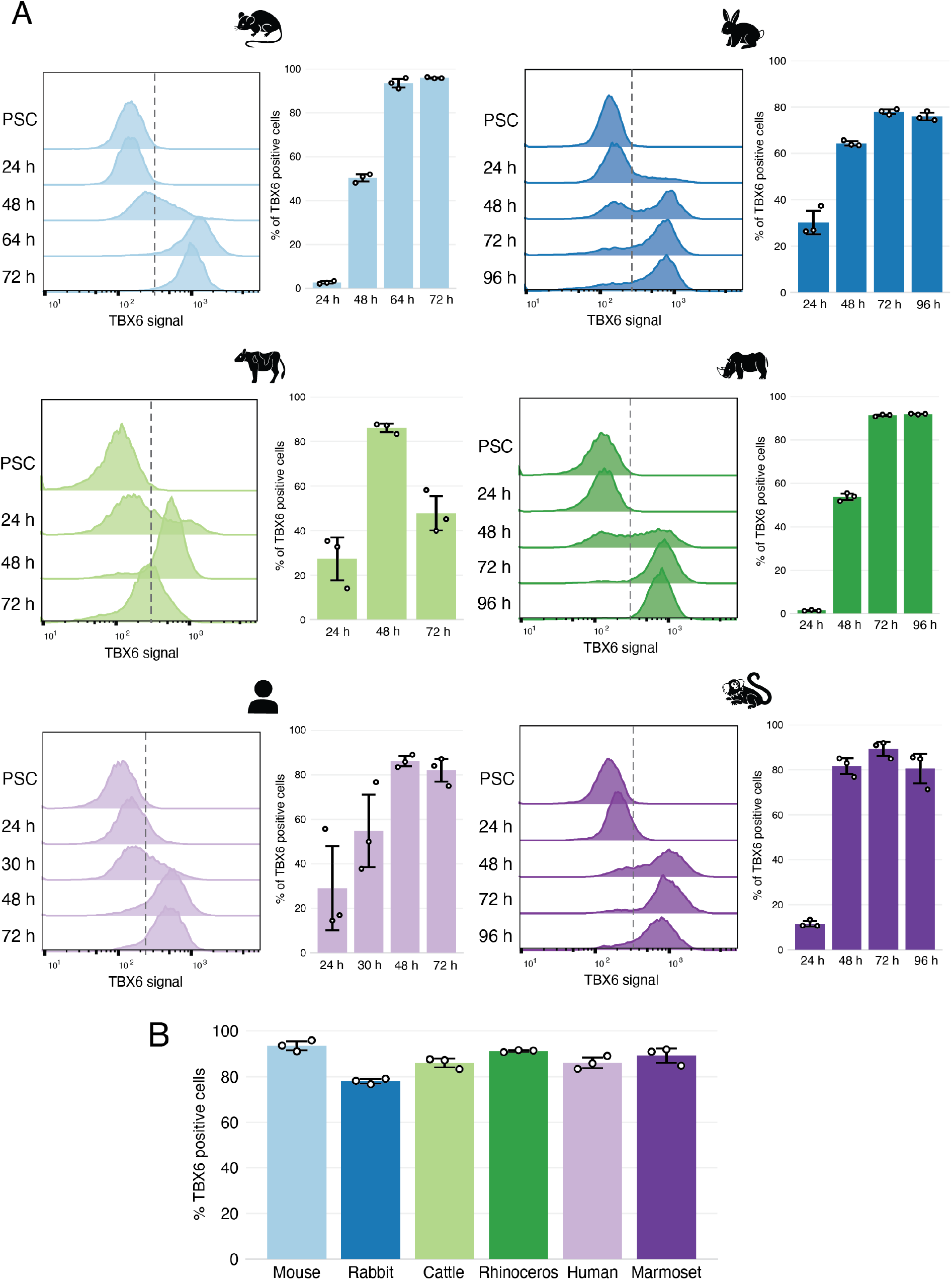
PSM differentiation efficiency across species (A) Left: Representative histogram of flow cytometry analysis during the course of PSM induction for six species. Staining signal of a PSM marker TBX6 at each day of differentiation is shown. Dashed line represents the threshold used to determine positive cells compared to the PSCs control. Right: PSM induction efficiency over the course of differentiation for six species. Percentage of cells expressing TBX6 was assessed based on the threshold shown in the left (dashed line). Error bars indicate means ± SD (n= 3). (B) Maximum efficiency of differentiation estimated from (A). 72h for mouse, rabbit, rhinoceros and marmoset and 48h for cattle and human. Cells at this stage were used for further analyses. Error bars indicate means ± SD (n= 3).

**Figure S3:**
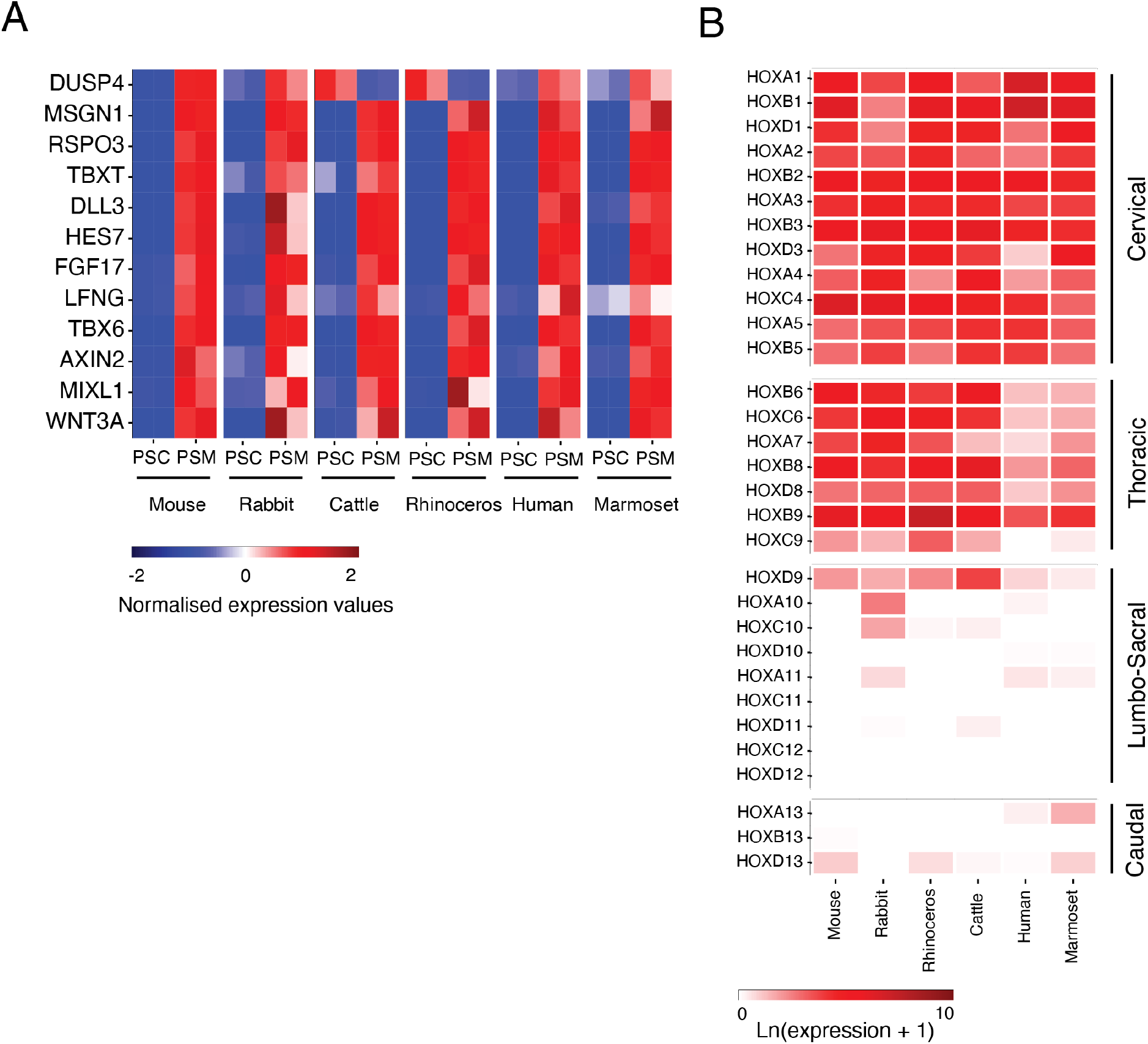
Expression of PSM markers and HOX genes in PSM-like cells (A) Heatmap of selected markers of PSM differentiation. Values for each gene were normalised to the mean of PSC and PSM samples of each species. (B) Heatmap of HOX gene expression levels in PSM cells of each species. Rows are the individual HOX genes ordered by anterior-posterior position. Anatomical anterior-posterior identity of each HOX group is indicated on the right.

**Figure S4:**
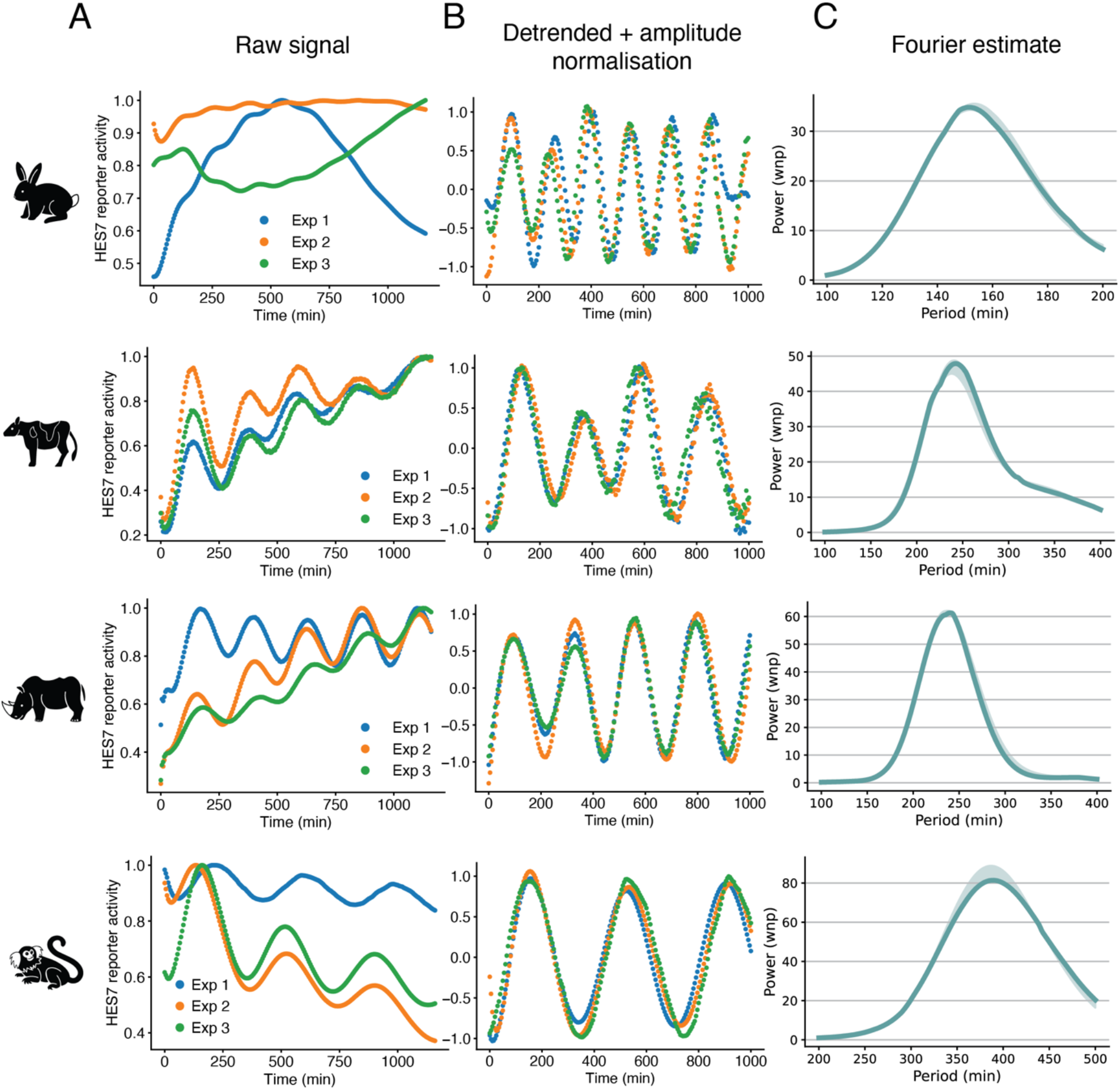
Quantification of the HES7 oscillatory period (A) Oscillatory HES7 reporter activity measured with a luminometer, using a collective signal from a 35-mm dish. Time course signals were normalised by the maximum signal, and results from 3 different experiments are shown. (B) Oscillatory HES7 reporter activity after detrending and amplitude normalisation using pyBOAT. (C) Fourier estimate of the processed oscillatory signals after wavelet analysis using pyBOAT. The period with the maximum power for each of the signals was used (Fig. 1E). Shading indicates median ± Q1/Q3 (n = 3).

**Figure S5:**
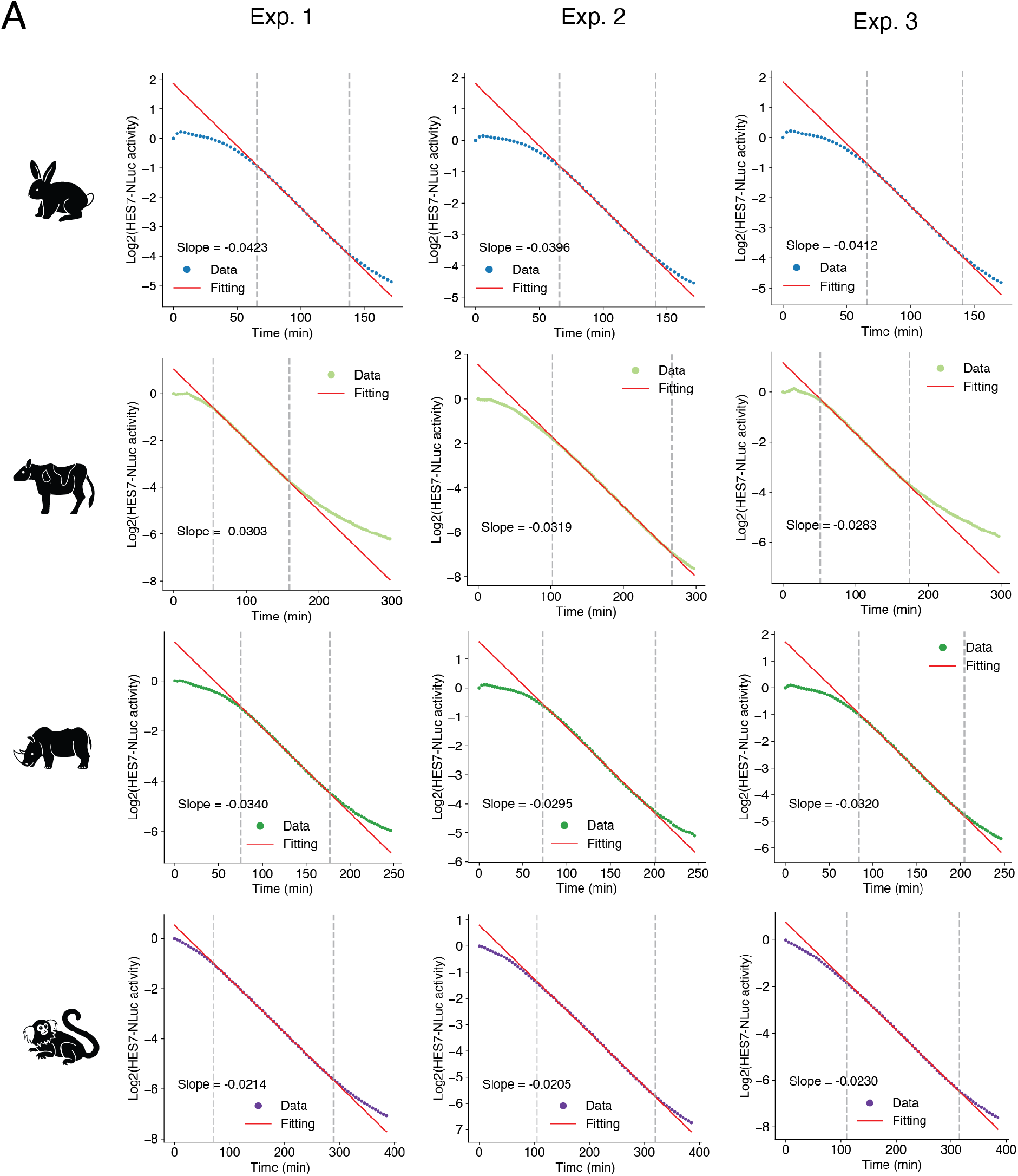
Quantification of the HES7 protein half-life (A) Fitting of HES7 protein degradation shown in Fig. 3B. Dashed lines indicate the most linear region considered by the RANSAC algorithm for the fitting. Slope of the fitted line is shown, and it was converted to the half-life using the equation: Half-life = -1/Slope.

**Figure S6:**
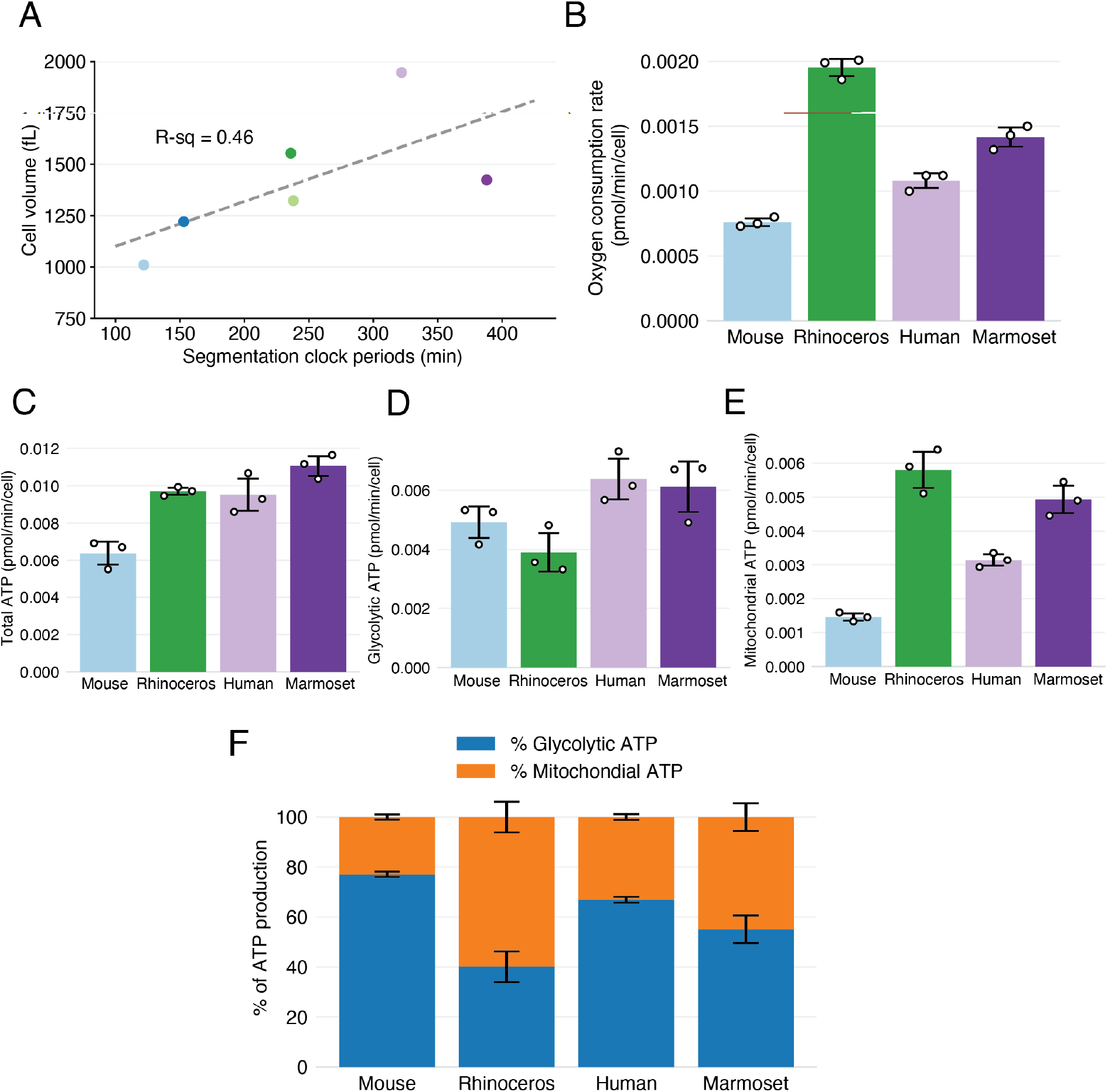
Cellular metabolic rates normalised by cell number (A) Scatterplot showing the relationship between the median cell volume and the segmentation clock period. Dashed line represents linear fitting. Colour scheme representing species is the same as Fig. 4A. R-squared value is shown. (B) Oxygen consumption rate per cell across species. (C) ATP production rate per cell across species. (D) Glycolytic rate of ATP production per cell across species. (E) Mitochondrial rate of ATP production per cell across species. (F) Ratio of glycolytic ATP production to mitochondrial ATP across species. (B-F) Error bars indicate means ± SD (n = 3 biological replicates). Data for (B) - (E) are the same as Fig. 4D - 4G but normalised only by the cell number (not by the cell volume).

**Figure S7:**
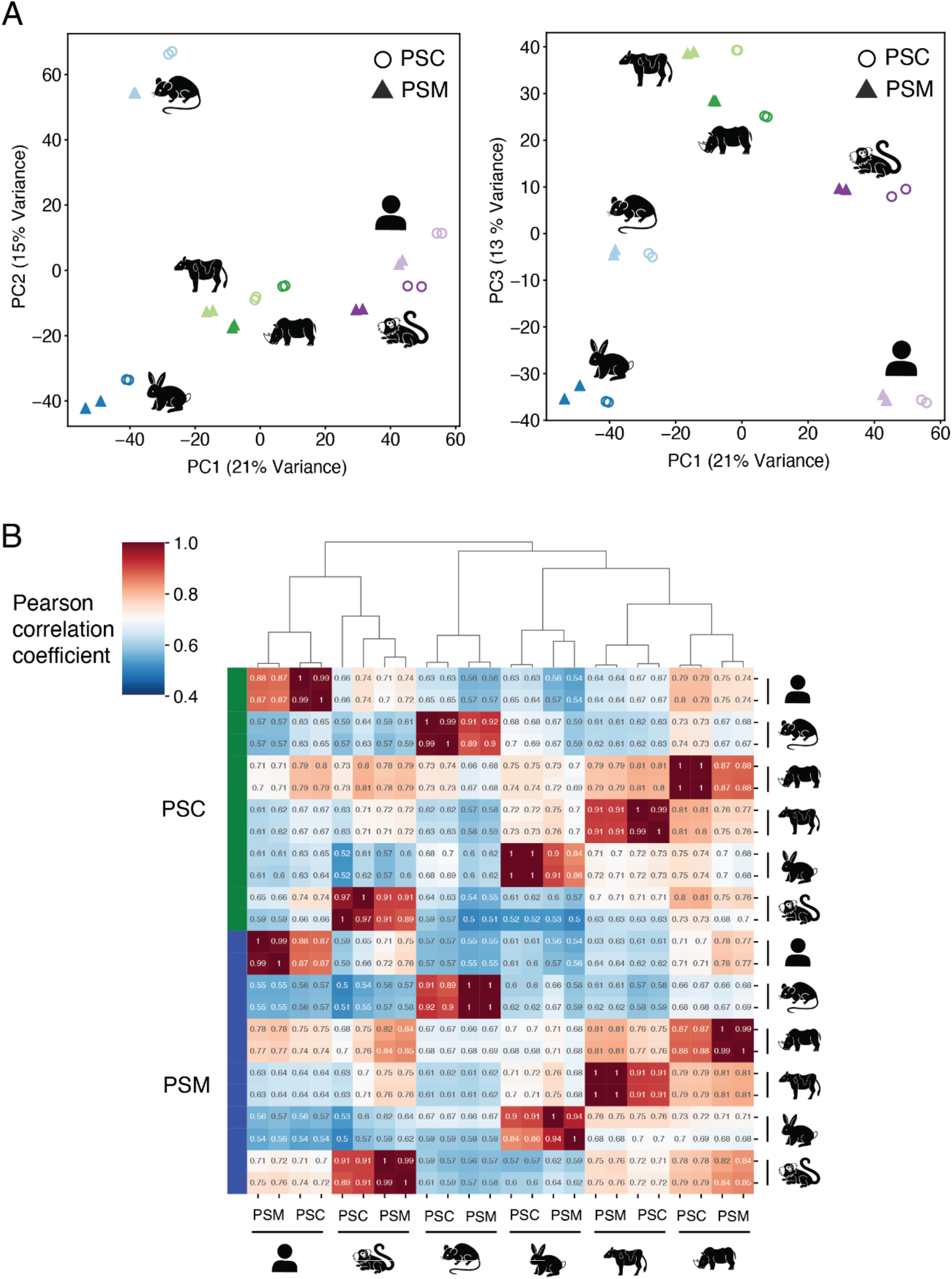
RNA-seq samples clustered by species instead of cell types (A) Principal component analysis (PCA) from bulk RNA-seq analyses. Two biological replicates of PSCs (circles) and PSM cells (triangles) of six species were used. Left: Components 1 and 2 and shown. Right: Components 1 and 3 and shown. The variance explained by each component is indicated. (B) Hierarchical clustering of all RNA-seq samples.

**Figure S8:**
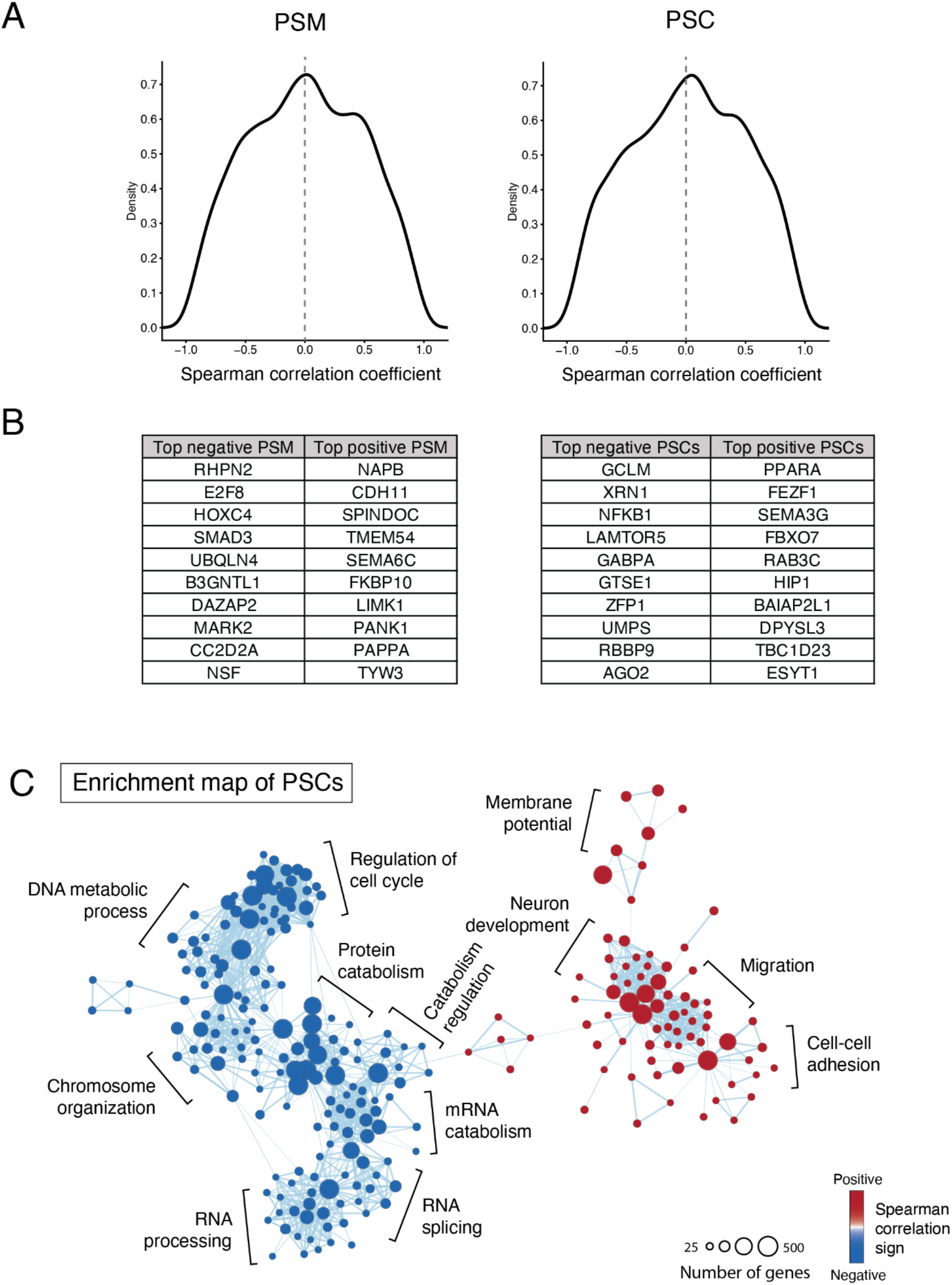
Correlation between gene expression and the segmentation clock period (A) Density plots showing the distribution of Spearman correlation coefficients between the gene expression level and the segmentation clock period. Results from PSM samples (left) and PSCs samples (right) of six species. (B) Top 10 genes that showed positive or negative spearman correlation coefficients. Results from PSM samples (left) and PSCs samples (right). (C) Enrichment map network of genes that showed correlated expression patterns with the segmentation clock period in PSC samples. Each dot represents a GO biological process functional term. Groups of inter-related functional terms tend to cluster together.

## Methods

### Pluripotent stem cell maintenance

All cells were cultured on a 5% CO2, 37°C, normoxic and humidified incubator. Media were changed every day in all cases.

Mouse EpiSCs (RIKEN BRC, AES0204) ^1^ were maintained on fibronectin-coated dishes with DMEM-F12 containing 15% Knockout Serum Replacement, Glutamax (2 mM), non-essential amino acids (0.1 mM), β-mercaptoethanol (0.1 mM), Activin A (20 ng/ml), bFGF (10 ng/ml) and IWR-1-endo (2.5 μM). Cells were passaged every two days using a 3 min incubation with accutase (Thermo Fisher Scientific). ROCK inhibitor Y-27632 (10 μM) was added to the media on the moment of passaging.

Marmoset iPSCs ^2^ were maintained on Geltrex (Thermo Fisher Scientific)-coated dishes with StemMACS iPS-Brew (Miltenyi) containing IWR1 (3 μM), CGP77675 (0.3 μM), AZD77675 (0.3 μM), CHIR99021 (0.5 μM), Forskolin (10 μM), Activin A (1 ng/mL) and OAC1 (1 μM). Cells were passaged every five days using a 5 min incubation in Versene (Gibco) followed by 5 min incubation in Collagenase IV (1 mg/mL). Pro-Survival compound (5 μM) was added to the media on the moment of passaging.

Rabbit ESCs (RIKEN BRC, AES0174) ^3^ were maintained on Matrigel (Corning)-coated dishes with the media composed of 50% mTESR1 (StemCell Technologies) and 50% DMEM-F12 containing 20% Knockout Serum Replacement, Glutamax (2 mM), non-essential amino acids (0.1 mM), β-mercaptoethanol (0.055 mM) and bFGF (10 ng/mL). Cells were passaged every two to three days using a 2 min incubation with accutase (Thermo Fisher Scientific). ROCK inhibitor Y-27632 (10 μM) was added to the media on the moment of passaging.

Human iPSCs (feederless 201B7, from RIKEN BRC HPS0063) ^4^ were maintained on Matrigel (Corning)-coated dishes or plates with StemFit medium (Ajinomoto). Cells were passaged every four days using a 3 min incubation with accutase (Thermo Fisher Scientific). ROCK inhibitor Y-27632 (10 μM) was added to the media on the moment of passaging.

Cattle ESCs ^5,6^ were maintained on Matrigel (Corning)-coated 35 mm dishes with mTESR1 (StemCell Technologies) or StemFit (Ajinomoto) base medium supplemented with bFGF (20 ng/mL), Activin A (20 ng/mL) and IWR1 (2.5 μM). Cells were cultured at high density and split every two days using 1-2 min TrypLE (Gibco) incubation. ROCK inhibitor Y-27632 (5 μM) was added to the medium in the moment of passaging.

Rhinoceros ESCs ^7^ were maintained on Matrigel (Corning)-coated dishes with the media composed of 50% mTESR1 (StemCell Technologies) and 50% DMEM-F12 containing 20% Knockout Serum Replacement, Glutamax (2 mM), non-essential amino acids (0.1 mM), β-mercaptoethanol (0.055 mM) and bFGF (10 ng/mL). Cells were passaged every four to five days using a 3 min incubation with 0.25 mM EDTA followed by dissociation into small clumps.

### Induction of PSM cells

Mouse EpiSCs were seeded on a 35mm dish coated with Matrigel and cultured in maintenance media without IWR-1 for one day. Then, mouse PSM was induced by culturing the cells for two days in CDMi ^8^ containing SB431542 (10 μM), DMH1 (2 μM), CHIR99021 (10 μM) and bFGF (20 ng/ml); the media will be hereafter referred to as SCDF media.

Rabbit, rhinoceros and cattle ESCs were seeded on a 35 mm dish coated with Matrigel. The next day, the media were changed to SCDF media, and cells were cultured for three more days in case of rabbit and rhinoceros and two more days in case of cattle.

Marmoset and human iPSCs were seeded on a 35mm dish coated with Matrigel and cultured for two to three days. Then the media were changed into CDMi containing bFGF (20 ng/ml), CHIR99021 (10 μM) and Activin A (20 ng/ml) for 24 hours. The human and marmoset cells were then further cultured in SCDF media for one and two days, respectively.

### TBX6 staining for flow cytometry

Cells were dissociated with accutase (Thermo Fisher Scientific) for 5 min at 37 °C and fixed in 4% paraformaldehyde in PBS for 10 min at room temperature (RT). For staining, 3 × 10^5^ cells were used. Cells were incubated overnight with an anti-TBX6 antibody (Abcam ab38883, 1:250) on Perm/Wash buffer (BD) at 4°C. The next day, cells were washed twice with Perm/Wash buffer and incubated with a 647nm Alexa Fluor secondary antibody (1:500) at RT for 2 hours. Cells were resuspended in 0.5 ml of Perm/Wash buffer and filtered for data acquisition on a LSRII cytometer (BD). Ten thousand events gated as single cells were recorded. Analysis was performed using FlowJo software. Stained PSCs were used as a control to set the intensity threshold. PSM cells with intensities over the threshold were considered as TBX6 positive.

### DNA constructs

The genetic constructs used in this study were obtained from Matsuda et al. 2020 ^9^. For the HES7 reporter construct (Fig. 1D), the hHES7 promoter and FLuc-NLS-PEST-UTR (hHES7) constructs were used. For the HES7 protein degradation construct (Fig. 3B), the rTetOne promoter and hHES7-NLuc (w/o intron) constructs were used. For the HES7 intron delay construct (Fig. 3C), the hHES7 promoter, NLuc-NLS-PEST-stop-hHES7 (w/o intron) and FLuc-NLS-PEST-stop-hHES7 (w/intron) constructs were used. The promoters or genes were subcloned into pDONR vector to create entry clones. These entry clones were recombined with a *piggyBac* vector (a gift from K. Woltjen) ^10^ by using the Multisite Gateway technology (Invitrogen). The constructs were stably introduced into the cells with by electroporation with Amaxa Nucleofector (Lonza) or using lipofectamine (Invitrogen) in the case of cattle cells.

### Oscillation analyses

After the induction of PSM, D-luciferin (200 μM) was added into the medium to monitor oscillations of the HES7 reporter signal. Bioluminescence was measured with Kronos Dio Luminometer (Atto). The obtained signal was analysed with pyBOAT 0.9.6 ^11^, a python-based software for time-frequency analysis of biological data. A threshold of 500 min was used for Sinc-detrending and amplitude normalisation of the signal in marmoset, cattle and rhinoceros cells. For rabbit cells, a 250 min threshold was used. The processed signal was then analysed using wavelets with a period ranging from 100 to 500 min. A Fourier estimate of the wavelet analysis provided a distribution of periods and its corresponding power. The period with the maximum power for each of the signals was considered. For plotting purposes, time-series data displayed were normalized to the first peak of oscillations.

### Organism characteristics

The approximate period of *in vivo* somite formation was calculated using studies describing the number of somites in staged embryos. Linear correlation between the embryonic day and the number of somites was used to extract the somite formation period. Note that these somite counts have a lot of uncertainty due to the difficulties in obtaining and accurately staging embryos from unconventional mammalian species. The values and references can be found on Table 1. Values of the average adult body weight and gestation length of the different species were obtained from the AnAge database (Build 14, visited on August 2022, https://genomics.senescence.info/species/index.html). The length of embryogenesis was extracted from different embryology manuals. The exact values and references can be found on Table 2.

**Table 2:**
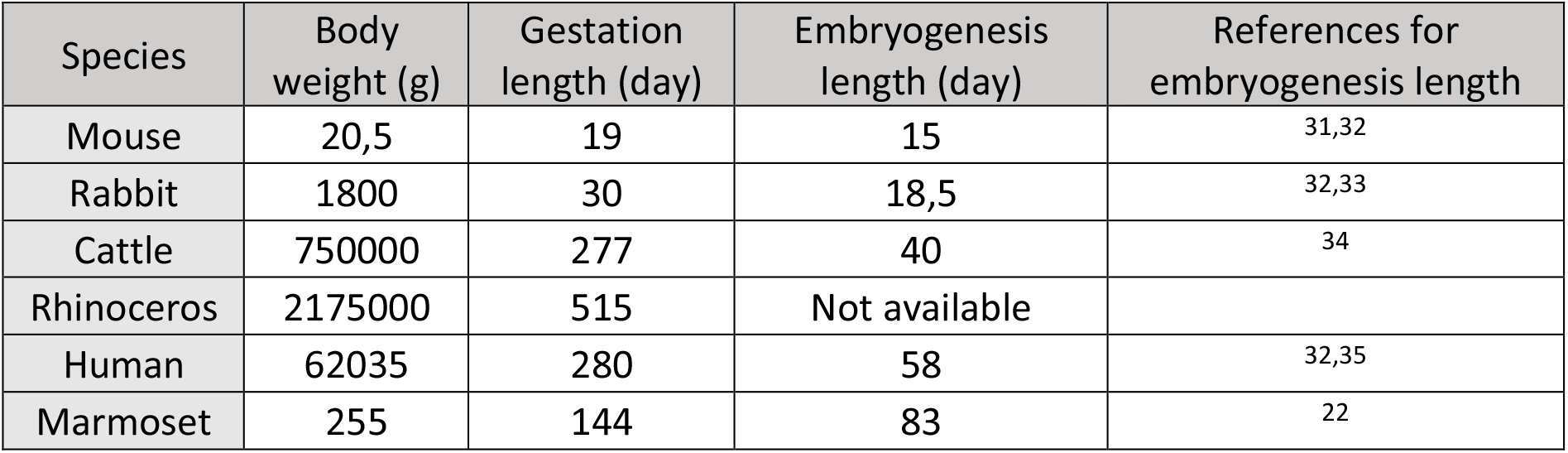
Organism parameters Average adult body weight and gestation length from the different species extracted from the AnAge database. Embryogenesis lengths extracted from different embryology manuals.

| Species | Body weight (g) | Gestation length (day) | Embryogenesis length (day) | References for embryogenesis length |
| --- | --- | --- | --- | --- |
| Mouse | 20,5 | 19 | 15 | 31,32 |
| Rabbit | 1800 | 30 | 18,5 | 32,33 |
| Cattle | 750000 | 277 | 40 | 34 |
| Rhinoceros | 2175000 | 515 | Not available |  |
| Human | 62035 | 280 | 58 | 32,35 |
| Marmoset | 255 | 144 | 83 | 22 |

### Phylogenetic tree reconstruction

The phylogenetic tree relating the six species was obtained by subsetting the mammalian tree published by Upham et al. 2019 ^12^ using their online tool (http://vertlife.org/phylosubsets/).

### HES7 protein degradation assay

As described in Matsuda et al. 2020 ^9^, the overexpression of a fusion construct of HES7 and NLuc was regulated by the rTetOne system (reverse TetOne system). After PSM cells were induced in the presence of Doxycycline (Dox; 100 ng/ml), the expression of the fusion protein was initiated by washing out Dox and changing the media into CDMi containing protected furimazine (Promega; 1 μM). After the NLuc signal was confirmed 5-8 hours later, the expression of the fusion protein was halted by Dox (300 ng/ml) addition, and the decay of NLuc signal was monitored with Kronos Dio luminometer. To exclude the influence of residual mRNAs, only the later time points where the NLuc signal displayed a single exponential decay curve were considered. To estimate the protein half-life of HES7, the slope of log2-transformed data was calculated. A RANSAC algorithm (scikit-learn) was used to find the most linear part of the decay curve.

### HES7 intron delay assay

As described in Matsuda et al. 2020 ^9^, the HES7 promoter-NLuc-stop-HES7 (w/o intron) and HES7 promoter-FLuc-stop-HES7 (w/intron) reporter constructs were introduced into the PSCs. After PSM cells were induced, the media was changed into CDMi containing protected furimazine (1 μM) and D-luciferin (1 mM), and the oscillations of the NLuc and FLuc signals were simultaneously monitored with Kronos Dio luminometer. To estimate the intron delay of HES7, the oscillation phase difference between the ‘w/o intron’ and ‘w/intron’ reporters was estimated by calculating their cross correlation with python (SciPy). Differences in the maturation time between NLuc and Flu were assumed to be constant across species based on human and mouse data ^9^.

### Cell volume quantification

Cells were dissociated with accutase (Thermo Fisher Scientific) for 5 min at 37 °C and washed in IMDM media (Sigma) with an osmolarity of 293 mOsm. Volume was measured on a Z2 Coulter counter (Beckman). The measured range was set from 7 to 21 microns in diameter. During the measurement, the cells were maintained in IMDM media. Approximately 6 × 10^4^ - 8 × 10^4^ cells were measured per experiment. The cell volume distributions were analysed using python.

### Seahorse metabolic rate analysis

PSM cells were dissociated with accutase for 5 min at 37 °C during the most efficient day of differentiation and re-seeded into fibronectin-coated Seahorse plates (Agilent) at a density of 7.27 × 10^5^ cells per cm2 in 100 μL of Seahorse XF DMEM (Agilent) supplemented with 10 mM glucose (Agilent), 1mM pyruvate (Agilent) and 2 mM glutamine (Agilent). Cells were allowed to attach at RT for 15 min and then transferred to a 37°C incubator without CO_2_ for 40 min. After that time, 400 μL of Seahorse XF DMEM media at 37°C were added carefully to each well without disturbing the attached cells for a total of 500 μL. Cells were incubated at 37°C without CO_2_ for 15 more min The Seahorse cartridge was hydrated overnight. For the real-time ATP rate assay (Agilent), 1 μM oligomycin, 0.5 μM rotenone and 0.5 μM antimycin A were used. All samples were run in seven to ten technical replicates in a Seahorse XFe24 (Agilent). Three biological replicates were performed for each species. The Wave Desktop and online app provided by the manufacturer was used for analysis.

### RNA library preparation

RNA samples were extracted from cultured cells using the RNeasy Mini Kit (Qiagen) following the manufacturer’s instructions. On column DNase digestion was performed on all samples. Barcoded stranded mRNA-seq libraries were prepared from 300 ng of high-quality total RNA samples using the NEBNext Poly(A) mRNA Magnetic Isolation Module and NEBNext Ultra II Directional RNA Library Prep Kit for Illumina (New England Biolabs (NEB), Ipswich, MA, USA) implemented on the liquid handling robot Beckman i7. Obtained libraries that passed the QC step were pooled in equimolar amounts; 2.1 pM solution of this pool was loaded on the Illumina sequencer NextSeq 500 and sequenced uni-directionally, generating ∼150 million reads, each 150 bases long.

### Gene expression analyses of cell types across species

Primary processing of the RNA-seq data was performed in the Galaxy ^13^ platform using a workflow composed of the main following steps:

1. Read cleaning using *Trim Galore!* (Galaxy Version 0.6.3) with automatic adaptor detection, Trim low-quality ends from threshold: 20, Overlap with adapter sequence required to trim a sequence: 1, Maximum allowed error rate: 0.1, reads becoming shorter than 20 were discarded.
2. Read Mapping using STAR ^14^ (Galaxy Version 2.7.8a) with default single-end options. Reads were mapped to hg38 (*H. sapiens*), mm10 (*M. musculus*), bosTau9 (*B. taurus*), calJac4 (*C. jacchus*), OryCun2 (*O. cuniculus*) and CerSim1 (*C. simum simum*).
3. Read filtering using Filter SAM or BAM, output SAM or BAM files on FLAG MAPQ RG LN or by region (Galaxy Version 1.8) to only keep mapped reads with MAPQ > 19 (which eliminates multi-mapping reads).
4. Stand-specific read counts were summarized at the gene level using featureCounts (Galaxy Version 1.6.3) with the reverse stranded option. The GTF files provided by GENCODE/Ensembl (v39 for human and vM23 for mouse, Bos_taurus.ARS-UCD1.2.106.chr.gtf for Cattle and Oryctolagus_cuniculus.OryCun2.0.106.chr.gtf for Rabbit) and RefSeq-based GTF provided by UCSC (cerSim1.ncbiRefSeq.gtf for Rhinoceros and calJac4.ncbiRefSeq.gtf for Marmoset) were used across all analysis.
5. RNA-seq data quality was assessed using FastQC (Galaxy Version 0.72) at different steps of the workflow to check sequencing quality and monitor filtering step efficiency, Picard CollectRnaSeqMetrics (Galaxy Version 2.18.2.1) to check the alignment of RNA to various functional classes of loci in the genome; finally, read trimming and read mapping reports were compared across samples for consistency and detect potential outliers using MultiQC ^15^ (Galaxy Version 1.9).

Pairwise gene orthology tables between each species and human were exported using Ensembl BioMart. For the rhinoceros, we used the gene orthology table (mouse-human-rhino) provided by Masafumi Hayashi and Katsuhiko Hayashi. Detailed methods for the construction of the mouse-human-rhino orthology table will be published separately. We then assembled a stringent multi-species gene orthology table using the human genes as the glue and considering an orthology one to one relationship type only.

Only the genes showing one to one orthology across all species were kept for further analysis. Raw counts were normalised using the Gene length corrected trimmed mean of M-values (GeTMM) method for best intra- and intersample comparisons ^16^. Reads per kilobase (RPK) were calculated for each gene using the gene length provided in the GFT file from each species. TMM-normalization was performed in R using the edgeR package ^17^. For PCA and correlation with the segmentation clock period, genes with a GeTMM value of less than 10 in all species were discarded.

### Principal component analysis

Principal component analysis was performed using the python library scikit learn. GeTMM values were log normalised before the analysis.

### Gene set enrichment analysis

Gene set enrichment analysis (GSEA) ^18^ was performed using the 4.2.3 version of the GSEA desktop app for iOS. All genes were pre-ranked by the values of Spearman’s correlation coefficients between the expression level and the segmentation clock period across six species. The gene set Gene Ontology (GO) biological process v7.5.1 from MSigDB was used ^19^. Only those gene sets with a size of more than 15 genes and less than 800 genes were kept for further analysis. Network visualization of similar terms was performed with the EnrichmentMap plug-in for Cytoscape 3.9.1 ^20,21^. Only those GO terms or pathways with FDR < 0.1 and p-value < 0.005 were shown in the network plots.

### Sample definition

All data are biological replicates.

